# Mammary microvessels are sensitive to menstrual cycle sex hormones

**DOI:** 10.1101/2023.04.21.537664

**Authors:** Carmen Moccia, Marta Cherubini, Marina Fortea, Akinola Akinbote, Prasanna Padmanaban, Violeta Beltran Sastre, Kristina Haase

## Abstract

The mammary gland is a highly vascularized organ that is influenced by sex hormones including estrogen (E2) and progesterone (P4). Beyond whole-organism studies in rodents or 2D monocultures, hormonal interactions and their effects on the breast microvasculature remains largely understudied. Recent methods to generate 3D microvessels on-chip have enabled direct observation of complex vascular processes; however, these models often use non-tissue-specific cell types, such as HUVEC and fibroblasts from various sources. Here, novel mammary-specific microvessels are generated by co-culturing primary breast endothelial cells and fibroblasts under optimized culture conditions. These microvessels are mechano-sensitive (to interstitial flow) and require endothelial-stromal interactions to develop fully perfusable vessels. These mammary-specific microvessels are also responsive to exogenous stimulation by sex hormones. When treated with combined E2 and P4, corresponding to the four phases of the menstrual cycle (period, follicular, ovular, and luteal), vascular remodeling and barrier function are altered in a phase-dependent manner. The presence of high E2 (ovulation) promotes vascular growth and remodeling, corresponding to high depletion of proangiogenic factors, whereas high P4 concentrations (luteal) promote vascular regression. The effects of combined E2 and P4 hormones are not only dose-dependent but also tissue-specific, as is shown by similarly treating non-tissue-specific HUVEC microvessels.

## 1. Introduction

The microvasculature is essential for constructing physiologic organ-on-chip systems, as the presence of blood vessels is critical to maintaining viability at scales limited only by diffusion-based transport. Moreover, microvessels are necessary for systemic drug delivery, nutrient exchange and clearance from tissues, making them important in the context of any preclinical study. [1] Several research groups have developed various strategies to incorporate perfusable microvasculature to model human-relevant disease on-chip.[2-4] Vasculature is formed *in vitro* by cell patterning, self-assembly, or a combination of both strategies. [5] In most reports, human umbilical vein endothelial cells (HUVECs) are co-cultured with stromal cells from various sources (including the lung [6, 7] and skin [8, 9]). Protocols for the isolation of HUVECs are reported more than 45 years ago and still represent the most common source of endothelial cells to study vascular biology.[10] Despite their ease of isolation, culture, and potential to form tubules, HUVECs are fetal and may not represent the phenotypic response of adult-derived endothelial cells. Recently, there has been a shift in focus toward generating tissue-specific models (reviewed in [3]); however, there is still an unmet need for tissue-specific vascular models.

Vascular heterogeneity exists between different organs and tissues, where metabolic demands are supported by different types of microvasculature.[11] Although phenotypic vascular differences are known to exist across tissues, the intrinsic genetic and epigenetic signatures promoting specialized organ-specific vasculature are still poorly understood,[12, 13] but largely influenced by the microenvironment.[14] In particular, the presence of stromal cells is essential for vascular formation and maintenance.[4, 15] Several recent studies show the influence of tissue-specific fibroblasts on the transcriptome of endothelial cells, giving rise to different subtypes of cells with unique gene expression profiles.[16, 17] For this reason, the presence of organ-specific microvasculature is essential to build physiologically relevant platforms to understand the pathophysiology of an organ.

The mammary gland and breast tissue could greatly benefit from a model of mammary-specific vasculature, as breast cancer is the most commonly diagnosed cancer in women.[18] Tumor-on-chip systems of various designs have been developed to study breast cancer subtypes and related processes, as reviewed in [19, 20]. Yet, those that do include a functional microvasculature use HUVEC.[19] As the breast is a dynamic system and undergoes cyclic remodeling regulated by sex hormones during puberty, lactation, menopause, and even during the monthly menstrual cycle, we expect that microvasculature from this tissue may also be sensitive to hormones. Estrogen (E2) and progesterone (P4) are steroid hormones produced and released by the ovaries. These hormones direct the development of female sexual characteristics during puberty and ensure fertility. Once E2 and P4 bind their nuclear receptors in cells, they activate the transcription of a variety of target genes involved in proliferation, metabolism, cell signaling, and survival.[21, 22] This process leads to dynamic remodeling of the mammary gland which undergoes cycles of expansion and regression associated with changes in cell number, tissue composition, and architecture.[23]

The effect of E2 on the vascular system is multifactorial, as it modulates vascular function by stimulating vasodilation and vascular relaxation by stimulating the release of vasodilatory substances from the endothelium (prostacyclin and nitric oxide synthesis) as well as by decreasing the production of vasoconstrictor agents (cyclooxygenase-derived products, reactive oxygen species, angiotensin II, and endothelin-1).[24-27] Less is known about the effect of P4 on the circulatory system, which can have both vasoconstrictor and vasodilator effects depending on the location of the vessels and their level of exposure.[28] For example, it has been reported (in rats) that progesterone has a vasodilator effect on coronary arteries.[29] While in other studies, it is described as a vasoconstrictor during pregnancy (in rats).[30] Nevertheless, little is known about the effect of these hormones on organ-specific vasculature in humans since most studies use non-specific endothelial cells in 2D systems.[31-34]

Recognizing the importance of vascular heterogeneity, we developed a mammary-specific microvasculature in a macro-scale fluidic platform wherein we can control perfusion. This model is used to investigate the role of sex hormones in mammary vascular development and barrier function. By generating a tissue-specific mammary microvascular system, our results demonstrate that morphologic remodeling is promoted by fibroblast density and interstitial flow. In addition, for the first time, the role of combined sex hormones is shown in the developing breast microvasculature. By exogenous stimulation of E2 and P4, representing the four phases of the menstrual cycle, hormone sensitivity of breast microvasculature is revealed. These results demonstrate mammary microvasculature as a highly sensitive network that remodels in response to changes in fibroblast density, flow and hormones, which are all critical in the context of the pathophysiology of breast cancer.

## 2. Results

### 2.1. Interstitial flow promotes mammary microvascular connectivity and perfusion

Primary human mammary vascular endothelial cells (HMVECs) and human mammary fibroblasts (HMFs) sourced commercially were first characterized by flow cytometry to confirm the expected expression of known endothelial (CD31, VE-Cadherin, endoglin) and fibroblast-associated markers (Fibroblast specific protein-1, FSP1, and α smooth muscle actin, αSMA), respectively (Figure S 1 A-B). Next, HMVECs and HMFs were co-cultured in a macro-fluidic design previously reported and fabricated in-house.[35-37] Adapting our previously published protocol [35], cells were seeded in a fibrin hydrogel, with an endothelial to stromal cell ratio of 5:1, and after several days in culture formed a vascular network with HMFs strongly associated with vessels **(Figure 1 A)**. Although these initial experiments demonstrated the formation of microvessels, they were quite narrow and did not present perfusable vascular lumens **(Figure 1 A and Bii)**.

**Figure 1.**
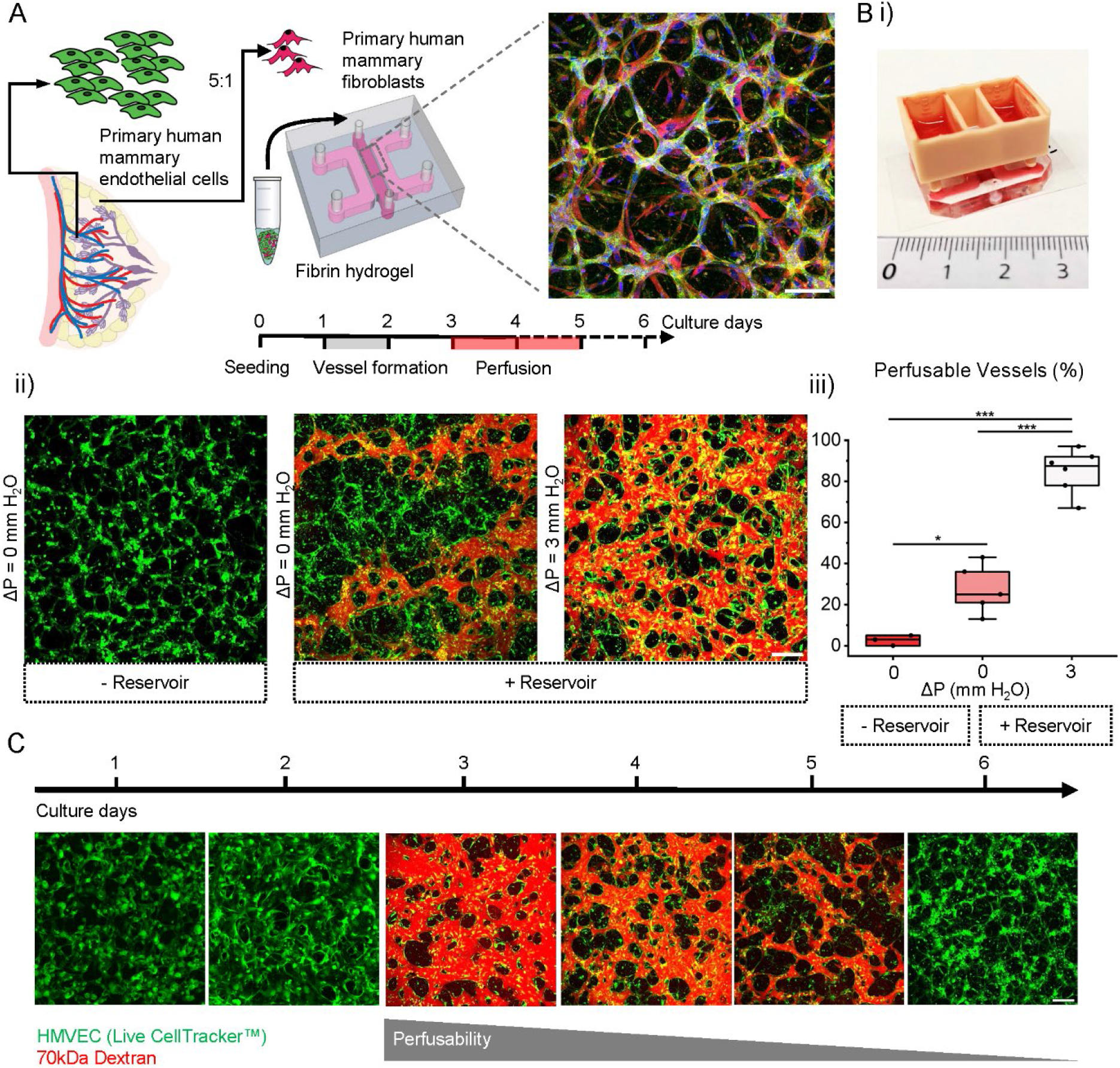
A (left) Graphical representation of the procedure used to generate mammary microvessels. Representative confocal maximum projection of fixed and stained microvessels at the fourth day in culture. Microvessels stained with endothelial marker CD31 green, fibroblast-specific protein-1 (FSP-1, red) and nuclei with DAPI (blue). Scale bar is 100 μm. B) I) Custom-made 3D printed reservoir for producing interstitial flow. II) Representative confocal images of microvascular network generated at different conditions: static culture without a reservoir, and with a reservoir under static (ΔP=0 mm H2O) and flow conditions (ΔP=3 mm H2O). HMVEC in the system are visualized by CellTracker™ green and the vessels perfused with Texas Red 70 kDa dextran. Scale bar is 200 μm. III) Percentage of perfused vessels in all three culture conditions for N=2 biological repeats. Box plots demonstrate median, percentile 25-75 quartile (box edge) and 10-90 (outer whiskers). Significance is shown by *p < 0.05 and ***p < 0.001 using one-way ANOVA with Tukey means comparison test. C) Representative confocal images of microvascular network at different days in culture. HMVEC in the system are shown by CellTracker™ green and perfusable vessels by Texas Red 70 kDa dextran. Scale bar is 200 μm. The grey bar indicates the lack of perfusion over time of the vessels.

Interstitial flow has been shown to promote early vessel formation.[19, 38] Hence, we employed fluidic reservoirs from day 1 to establish an intermittent interstitial flow, which was re-established daily. A range of pressure gradients (0-7 mm H2O) was tested. This applied interstitial flow resulted in a significant increase in perfusable vessels, around ∼85%, demonstrated by dextran perfusion compared to the static condition (0 mm H2O), with only ∼28% perfusable vessels with the same volume of media **(Figure 1 Biii)**. Increasing the pressure gradient (from 3, 5 and to 7 mm H2O) demonstrated a significant increase in vessel diameter and changes in other morphologic characteristics. However, the most physiologic vessels were generated using 3 mm H2O (Figure S 2 B-D).

Under this pressure gradient (3 mm H2O used for subsequent studies), mammary microvessels form rapidly—with branched and connected vascular networks formed in less than 24 hours from initial seeding **(Figure 1 C)**. This behavior contrasts other previously reported microvessels cultured from either HUVEC or iPSC-ECs in the same device which form between 5-7 days in culture.[36, 39] Notably, although these mammary microvessels form rapidly, they begin to regress after 4 days in culture.

### 2.2. Mammary fibroblasts affect microvascular morphology

It is widely recognized that fibroblasts are critical for supporting angiogenesis.[36, 39, 40] Fibroblasts in the breast are known to exist in lobular and interlobular positions in the mammary gland; however, little information exists about their role in vascular development and maintenance.[41] Thus, HMVECs and HMFs were co-cultured at different cell ratios 5:1,10:1 and 20:1, while keeping the final concentration of endothelial cells constant (6×10^6^ cells/mL) **(Figure 2 A)**. By varying the total number of fibroblasts, a striking change in vascular morphology was observed **(Figure 2 A-D)**. An increased fibroblast concentration led to a significant decrease in microvessel diameter. Microvessels co-cultured at the 5:1 ratio resulted in a mean vessel diameter of 30 ± 20 µm, comparable to *in vivo* values for adult women (15-50 μm).[42, 43] The mean branch length of mammary vessels remained unchanged with varied fibroblast concentrations, yet branch density was significantly increased with increasing fibroblasts (5:1 ratio) **(Figure 2 C-D)**. Interestingly, the number of fibroblasts did not have any measurable effect on vascular barrier function (permeability to 70 kDa dextran), which was comparable among the different ratios **(Figure 2 E)**.

**Figure 2.**
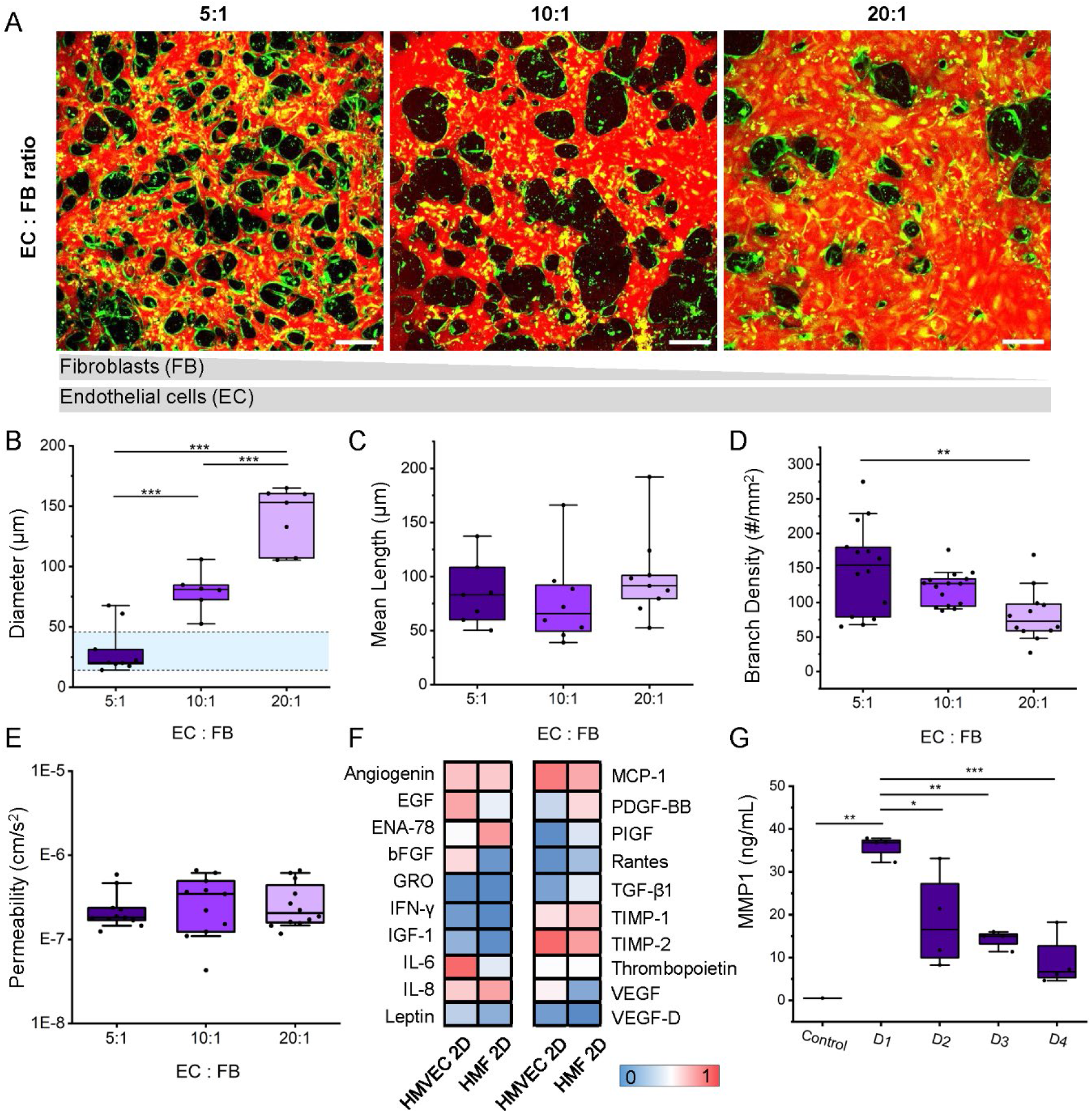
A) Confocal maximum projection images highlight microvessel formation at varied endothelial to fibroblast cell ratios. The grey scales represent the amount of fibroblasts and endothelial cells in the different conditions. HMVEC are shown with CellTracker™ green and microvessels are perfused with Texas Red 70 kDa dextran at day 4 to assess vessel perfusion. Scale bar is 200 μm. Morphologic comparison of the microvessels at the different ratios: B) effective diameter (blue region shown demonstrates average ex vivo mammary-specific measurements), C) mean length, D) branches density and E) permeability to 70 kDa dextran all for N=2 biological repeats. F) Semi-quantitative measure using a cytokine array of supernatant collected from HMVEC and fibroblast (HMF) cells in monolayer culture. G) MMP1 quantification by ELISA of control media (VL 5%) and supernatant from devices (5:1 ratio) from day 1 to 4. Box plots demonstrate median, percentile 25-75 quartile (box edge) and 10-90 (outer whiskers). Significance is shown by *p < 0.05, **p < 0.01, ***p < 0.001, for data following normality with one-way ANOVA with Tukey means comparison test, and if normality is rejected using Kruskal-Wallis ANOVA test.

We established that fibroblasts were necessary to form and maintain a functional mammary microvasculature. Without the fibroblasts in co-culture, thin yet connected vascular networks arise but regress rapidly after a few days in culture (Figure S 1 C). Fibroblasts are known to secrete crucial factors like vascular endothelial growth factor (VEGF), transforming growth factor-β (TGF-β), and platelet-derived growth factor (PDGF), that promote vessel growth.[44] Therefore, we performed a semi-quantitative analysis of angiogenic factors assessed for both HMVEC and HMF in monocultures **(Figure 2 F)**. Surprisingly, known angiogenic factors are expressed at relatively low levels in HMF (VEGF, basic fibroblast growth factor (bFGF), and TGF-β), but inflammatory cytokines (interleukins IL-6 and IL-8, monocyte chemoattractant protein-1 (MCP-1) and epithelial-neutrophil activating peptide (ENA-78)) were expressed at relatively high levels. Moreover, we also measured MMP1, an interstitial collagenase capable of degrading collagen types I, II, and III, which is also involved in vascular remodeling.[45] MMP1 was measured from supernatant collected from the 3D microvessels grown using the 5:1 ratio at different time points. MMP1 production significantly decreases over time supporting the strong remodeling observed on the first day following seeding **(Figure 2 G)**.

### 2.3. Sex hormones alter mammary microvessel morphology

The effects of 17β-estradiol and progesterone were investigated on mammary vascular development. First, the expression of E2 and P4 receptors was confirmed in both HMVEC and HMF by immunofluorescence (Figure S 3 A). Next, we generated mammary microvessels and supplemented normal media by daily perfusion of hormones at increasing concentrations (0, 1, 10, and 100 nM for E2, and 0, 1, 5, and 25 nM for P4) (Figure S 3 B). A dose-dependent effect was observed on vascular development with high concentrations leading to non-perfusable vessels. Since E2 and P4 act in unison and are reported at much lower plasma concentrations *in vivo* [46], we next examined the influence of these hormones combined as they vary across the menstrual cycle. Concentrations were chosen to mimic the mean concentrations of phases of the menstrual cycle: follicular, ovulation, luteal phase, and period **(Figure 3 A-B)**. Hormone treatments were exogenously added to the media the day after seeding (maintaining an intermittent interstitial flow, as before) and refreshed daily for 4 days. Treatment resulted in distinct morphologic changes, particularly when high levels of either estrogen or progesterone are present, as in the ovulation and luteal phases, respectively. This is demonstrated by CD31 staining of the microvessels **(Figure 3 A)**.

**Figure 3.**
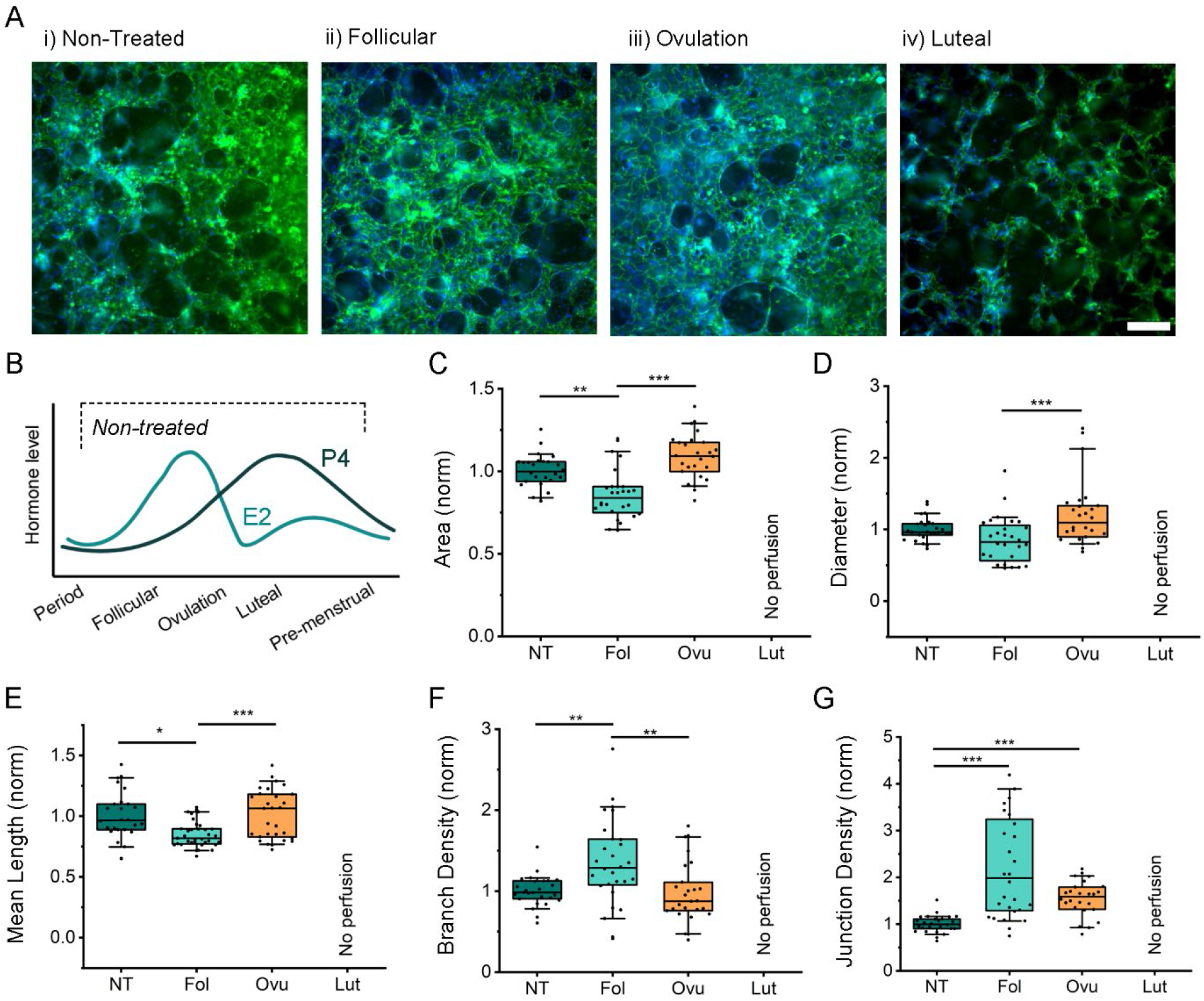
A) Confocal maximum projection images of microvessels at day 4 with different hormonal treatments mimicking the phases of the menstrual cycle. Microvessels stained with CD31 green and nuclei with DAPI (blue). Scale bar is 200 μm. B) Graphical representation of hormone oscillation during menstrual cycle. C-G) Comparison of morphological parameters of microvessels between the different hormonal conditions: period/non-treated, NT; Follicular, Fol; Ovulation, Ovu; Luteal, Lut. Normalized (norm) data from 5 separate experiments with ≥3 devices per condition. Shown are C) vessel area coverage, D) effective vessel diameter, E) mean branch length, F) branch density and G) junction density. Box plots demonstrate median, percentile 25-75 quartile (box edge) and 10-90 (outer whiskers). Significance is shown by *p < 0.05, **p < 0.01, ***p < 0.001 using Kruskal-Wallis ANOVA

To quantify vessel segments with connected and open lumens, FITC dextran was perfused in the mammary microvessels. Morphologic parameters were quantified on day 4, using ImageJ macros as described in our previous work. [1, 39] Considering that variability between biological repeats of mammary microvessels was observed, we compared hormone-treated samples to the non-treated (normalized) control for each experiment. Both the follicular and ovulation hormone treatments resulted in significant changes in the vascular area, with the ovulation concentrations leading to an increase in the vessel area coverage. The high E2 and low P4 associated with the ovulation phase also resulted in significant increases in vessel diameter **(Figure 3 D)** These treatments led to significant changes in branch and junction densities **(Figure 3 E-F)**. All mammary microvessels treated by the luteal concentrations (highest P4) demonstrated disconnected vasculature, as shown after 4 days and thus there were no perfused vessel segments to measure with our automated segmentation pipeline **(Figure 3 B)**.

Considering that E2 and P4 can both lead to cell proliferation in certain tissues,[47-49] we treated HMVEC and HMF (in 2D) with concentrations representing the different menstrual phases. After four days, HMVEC endothelial cells did not respond to treatments, unlike HUVEC, as shown by EDU assays (Figure S 4). However, HMF proliferation was significantly increased compared to non-treated cells.

### 2.4. Sex hormones alter mammary microvessel barrier function

In addition to morphological changes observed during vessel formation, we evaluated vascular endothelial barrier function in response to sex hormones. We performed time-lapse measurements to track the flux of 70 kDa dextran across the endothelial barrier—as a proxy for endothelial permeability to solutes similar in size (such as albumin) **(Figure 4 A)**. Since the luteal phase was not perfusable, measurements were evaluated only for the other defined phases. The solute permeability of these mammary vessels was comparable across treated and non-treated conditions. Although statistically insignificant, there was an increased trend in leakiness in the ovulation phase, which had the highest amount of 17β-estradiol **(Figure 4 B)**. The effect of hormones on cytokine production in these microvessels was also assessed by cytokine array (Figure S 5). Pooled supernatants were collected from 5 devices treated with the different hormonal conditions on day 4. Although semi-quantitative, results indicate a change across the different phases in proangiogenic factors including VEGF, placental growth factor (PLGF), bFGF, and epidermal growth factor (EGF). For this reason, we further investigated these proangiogenic factors by ELISA **(Figure 4 C-F)**. Supernatant from 4 devices for the different hormonal conditions was collected on day 2 (N=3 biological repeats), prior to any vascular regression. VEGF, which is supplemented in the media, was depleted equally between the different conditions. EGF and FGF, which are also present in the media, were differentially depleted between conditions. Not surprisingly, the follicular phase with increased vascular remodeling had the most significant depletion of these proangiogenic factors. Interestingly, PLGF, which is not present in the culture media, is expressed at high levels in the non-treated vessels as opposed to hormone-treated conditions.

**Figure 4.**
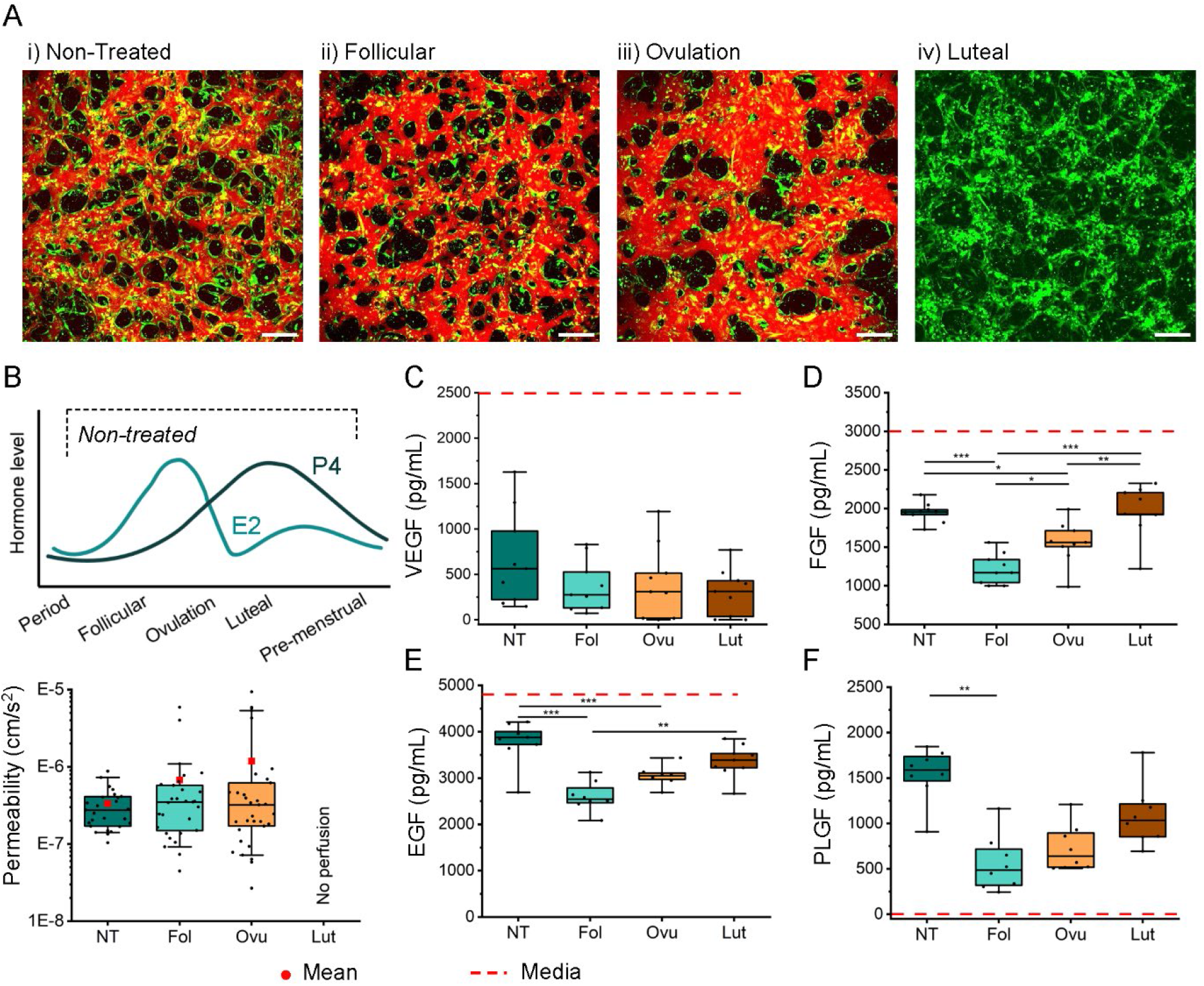
A) Confocal maximum projection images of mammary microvessels formed under different hormonal conditions at day 4. HMVEC shown by CellTracker™ green and microvessels perfused with Texas Red 70 kDa dextran at day 4 to assess vessel permeability. Scale bar is 200 μm. B) Vessel permeability to 70 kDa dextran is shown for the different hormonal conditions (non-treated, NT; Follicular, Fol; Ovulation, Ovu; Luteal, Lut). Data from 5 separate experiments with ≥3 devices per condition. ELISA assay performed from supernatant of devices collected at day 2 for C) VEGF, D) FGF, E) EGF and F) PLGF. Data is from 3 biological repeats with 3 devices per condition each. The dashed red lines in the plots indicates the level of each factors present in the control media. Box plots demonstrate median, red dots for the mean, percentile 25-75 quartile (box edge) and 10-90 (outer whiskers). Significance is shown by *p < 0.05, **p < 0.01, ***p < 0.001, using one-way ANOVA with Tukey means comparison test for data following normality, or if normality is rejected using Kruskal-Wallis ANOVA test.

### 2.5 The effect of sex hormones is endothelial cell-specific

Considering the observed hormonal sensitivity of mammary microvessels, we finally explored the effects of E2 and P4 on HUVEC microvessels. First, the expression of ER and PGR was determined by qPCR for HMVEC, HMF and HUVEC **(Figure 5 A)**. HUVEC shows a much higher expression of ER compared to the mammary endothelial cells, yet surprisingly HMFs express the highest expression for all receptors. Next, we generated vessels by co-culturing HUVEC and mammary fibroblasts using the same protocol as for the mammary-specific vessels **(Figure 5 B)**. Hormonal treatment for HUVEC-HMF co-cultures lasted for 7 days following seeding, since the vessels form between days 4-5. Non-treated vessels resulted in thin and non-perfusable networks **(Figure 5 Ai)**. The presence of hormones drastically changes the morphology of these co-cultured vessels, with a clear difference in area coverage and vessel diameter (Figure S 7). The presence of hormones promoted the generation of perfusable vessels. However, higher estrogen concentrations, as in the follicular and the ovular phases, resulted in very leaky vasculature, preventing permeability analysis **(Figure 5 A)**. Morphological parameters of HUVEC co-cultured vessels were normalized to NT samples and compared to HMVEC co-cultured vessels **(Figure 5 C-F)**. The comparison shows a clear difference in the diameter and area between the two co-cultured microvessels as they are treated with hormones.

**Figure 5.**
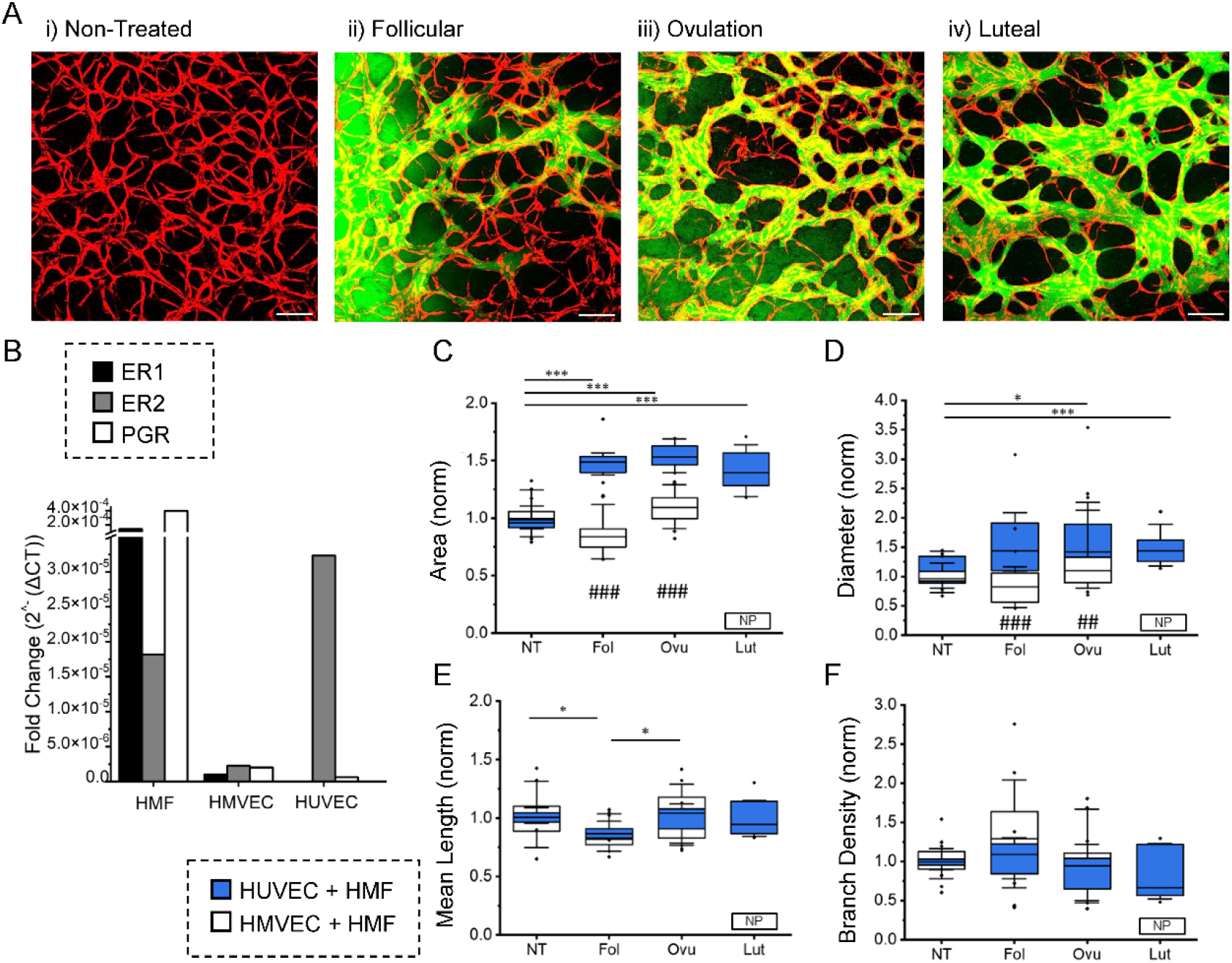
A) Confocal maximum projection images of mCherry-expressing HUVEC (red) and HMF (non-fluorescent) co-cultured microvessels formed under different hormonal conditions (non-treated, NT; Follicular, Fol; Ovulation, Ovu; Luteal, Lut). Microvessels are perfused with FITC 70 kDa dextran green on day 7 to assess vessel permeability. Scale bar is 200 μm. B) Genetic expression of ER1, ER2 and PGR in HMF, HMVEC and HUVEC grown in monocultures. (C-F) Morphological comparison of the vessel from HUVEC co-cultures (blue) and HMVEC co-cultured vessels exposed to hormone treatments. Shown are C) vessel area coverage, D) effective diameter, E) branch length and F) branch density. NP = No perfusion. Box plots demonstrate median, percentile 25-75 quartile (box edge) and 10-90 (outer whiskers). N=3 biological repeats for HUVEC co-cultures. Significance is shown by *p < 0.05,, **p < 0.01, ***p < 0.001 for HUVEC co-cultured vessels (across treatments), while significance between vessel types is shown by ## p < 0.01, ### p < 0.001 using one-way ANOVA with Tukey means comparison test.

## 3. Discussion

The generation of blood vessels *in vitro* is crucial for mimicking *in vivo* organ and tissue function, as blood vessels are essential for characteristic tissue growth, properties, and maintenance. Several organ-on-chip models have successfully integrated a vascular network, with remarkable success in the lung, liver, skin, blood-brain barrier, and kidney.[50-54] One of the limitations of vascularized organ-on-chip technology is the use of HUVEC and other non-specific endothelial cells.[55] The importance of endothelial cell heterogeneity in terms of phenotype, gene expression, antigen composition, and their specific function has been recently highlighted.[13] For this reason, the presence of tissue-specific cells in vascularized organ-on-chip technologies is the next step toward a system that closely resembles the *in vivo* physiology of human organs.[19] Little is known about the vasculature in the dynamic mammary gland, thus, for the first time, perfusable mammary microvascular networks with breast-specific cells were generated.

Herein, HMVEC-HMF co-cultures were optimized to generate perfusable mammary microvessels. These vessels demonstrate rapid vascularization, characterized by the appearance of an extensively branched and connected endothelial network as early as 24 hours post-seeding, with full perfusion by days 3-4. In contrast, previously reported vascular networks derived from other co-cultures achieve perfusion after nearly one week in culture.[36, 39] In the mammary system, accumulation of cells in the vessel lumen and subsequent negative remodeling of vessels occurs by day 5 **(Figure 1 C)** limiting long-term *in vitro* studies. Our co-cultures are generated from the only commercially available human mammary endothelial cells and fibroblasts. Although generating mammary microvessels from other donors would be of significant interest, isolating breast endothelial cells from reduction mammaplasties has a very low success rate and they do not typically survive in culture.[56, 57]

The pressure gradient, driven by a beating heart, reduces along the branched circulatory network towards microvessels – this flow is necessary for the delivery of oxygenated blood and immune cells, and nutrient delivery. Changes in vascular dynamics are well-known to affect the behavior of endothelial cells, which has been recapitulated *in vitro*.[58] Here, gravity-driven flow was established in our microvessels through a 3D printed reservoir, as an interstitial flow has been also previously shown to induce early vessel formation. Mammary microvessels in our system are highly mechano-sensitive, with higher pressure gradients resulting in significant morphological and barrier function changes (Figure S 2). At a pressure gradient equal to 3 mm H2O, vessels achieved a physiologic diameter range (mean 30 ± 20 μm) within the range of 15-50 μm reported in humans.[43] Heterogeneity in branch patterns occurred near regions of increased interstitial shear stress (as shown by a computational fluid dynamics (CFD) simulation of the device, Figure S 8). These microvessels demonstrate similar behavior to placental-like HUVEC vessels, where an applied interstitial flow was shown to promote vessel formation, diameter enlargement, and reduced permeability to solutes.[58]

Stromal cells secrete cytokines, growth factors, and proteases that act as angiocrine signals on endothelial cells, stimulating their activation, promoting vascular remodeling, and influencing function.[4, 15] Previous research from our group has shown that tissue-specific stromal cells modulate vascular morphology and barrier function, highlighting the need for the use of organ-specific cells.[39] Therefore, we hypothesized that mammary fibroblasts are important contributors to mammary vessel formation. Notably, breast endothelial cells cultured alone show discontinuous vasculature in the absence of fibroblasts (Figure S 1-C). The presence of fibroblasts allows the formation of vasculature and there is a strong correlation between fibroblast density and vascular morphology. A high fibroblast concentration results in smaller vessel diameters resembling those seen in biopsies *in vivo*.[43] MMP1, a metalloprotease that degrades collagen type 1, is strongly expressed in the first days of co-culture, indicating that matrix degradation occurs during the earliest stages of network formation. The mammary gland is rich in stromal tissue and fibroblasts are the predominant stromal cell type. Under physiological conditions, fibroblasts are essential in the mammary gland and change during the different hormonal phases.[59] During pregnancy and lactation, for example, fibroblasts regulate the composition, organization, and stiffness of the extracellular matrix (ECM), which controls epithelial cell proliferation, migration, differentiation, and polarity, facilitating the formation of new lactiferous alveoli and supporting blood microvessels.[59] Fibroblast function is strongly influenced by sex hormones herein, as shown by the fact that mammary fibroblasts express higher levels of estrogen and progesterone receptors than endothelial cells. Nevertheless, the effects of sex hormones on breast fibroblasts remain poorly understood, as most studies on these cells have focused on their role in breast cancer, in which, cancer-associated fibroblasts (CAFs) are the most studied. CAFs represent the predominant cell population (∼80%) in the breast tumor microenvironment and are involved in breast cancer development, progression, and metastasis.[60] While outside the full scope of this study, investigating the contributions of breast fibroblasts in normal physiological function will help us better understand their role in regulating the breast microvasculature.

Hormonal oscillations during the menstrual cycle affect the mammary epithelium, ECM, stromal compartment, immune system, and vascular network.[61] Previous studies on hormone interactions with endothelial cells in 2D reported an increase in proliferation and angiogenesis in presence of estrogen.[62-64] Additionally, alterations in vascular density within the mammary gland have been documented during the phases of pregnancy and lactation, suggesting a direct influence of sex hormones concentration on vascular growth and remodeling.[65] In order to shed light on the effects of E2 and P4 on vascularization and endothelial barrier function, we perfused our mammary-specific microvessels with hormone concentrations corresponding to the different phases of the menstrual cycle, as reported in.[66] Although hormone concentrations fluctuate during the menstrual cycle, for practical reasons (a practical limitation here) we chose an average value representing each phase. After 4 days of hormone treatment, striking changes in vascular morphology were observed **(Figure 3 A)**. In particular, hormones that mimic the luteal phase (high P4) resulted in progressive regression of vessels. One explanation for a reduction in vascular density could be due to enhanced proliferation and competition of HMFs, which are proliferative in response to these treatments in 2D **(Figure S 4 B)**. Significant vascular remodeling was also observed in the follicular phase

-decreased area and diameter and increased branching and junction densities. This result could be explained by E2 peaks in the follicular and luteal phases, which are recognized as highly mitotic phases in the breast.[67] In fact, it is known that with increasing estrogen and progesterone, epithelial alveolar buds start to form and differentiate.[61, 68] In the ovulation phase (highest E2 concentration) significantly larger vessel areas and diameters were observed with reduced branch density. Despite the variability between experiments, a trend towards increased leakiness of the vessels can be observed during the ovulation phase. This is consistent with the function of estrogens in promoting vasodilation, increasing fluidic volume and permeability of the vessels.[27, 69] Analysis of four (VEGF, FGF, EGF and PLGF) important proangiogenic factors were also measured in response to hormone treatments. VEGF, which plays a key role in angiogenesis and is present in the vascular media, was depleted in all conditions (hormone treated or not) indicating the important role of this factor in the formation and maintenance of the microvasculature. FGF and EGF, two other proangiogenic factors were depleted mostly in the follicular phase, concomitant with increased vascular remodeling **(Figure 4 A-D)**. Moreover, PLGF, which is not present in the cell culture media, showed a similar behavior of depletion for the follicular phase. PLGF belongs to the VEGF family and promotes vascularization, therefore we hypothesize that in the follicular and ovulatory phases, with increased vascular remodeling, there is a negative feedback mechanism leading to reduced concentrations in comparison to untreated microvessels. Of course, the breast is much more complex than our reductionist model system, and other cell types (adipose, immune, epithelial and smooth muscle cells) are also influenced by hormones which affect the microvasculature. Additionally, other hormones such as follicle-stimulating hormones and luteinizing hormones also have an effect on the breast, this has not yet been fully elucidated and could be investigated in the future using our model.

We observed a clear effect of sex hormones on mammary microvessel in our model, however, whether this effect is mammary-specific was unclear. Thus, we also generated HUVEC-HMF co-cultures to examine the effect of hormones on varied endothelial cells. The differences between the two endothelial cultures treated are notable. For example, luteal treatment in the breast led to vessel regression after 4 days, whereas the same treatment in HUVEC microvessels results in a fully functional vascular network. One potential explanation for the observed difference is that HUVEC typically are exposed to high levels of P4 nearing term (although ∼ 1% of the total in maternal serum).[70], so it is possible this reinforces pro-angiogenic behavior in our system.

Vessel diameter increases under hormone treatment in HUVEC microvessels, while in the mammary microvessels, the diameter decreased in the follicular phase, but branch and junction density increase. One explanation for these differences could be found in the different expression levels of ER and PR in HUVEC and HMVEC **(Figure 5 A)**. Moreover, HMVEC cells originate from a single female donor, whereas HUVEC are from pooled (male and female) donors. Pooled donors may reduce donor-to-donor variability; however, we recognize the necessity of replicating experiments using endothelial cells from different donors. Our research underscores the significance of utilizing tissue-specific cells to investigate organ-specific characteristics, such as the function of sex hormones in breast microvasculature.

## 4. Conclusion

For the first time, perfusable primary mammary microvessels have been cultured on chip and used to demonstrate the effect of physiological concentrations of sex hormones. Estrogen and progesterone are shown to have a profound effect on breast-specific and non-specific microvessels at the menstrual level. This work highlights the importance of using tissue-specific cells and the role of sex hormones in breast vascular remodeling. In the future, this system will enable the study of pregnancy and menopause hormone levels in breast vessels. In addition, this system could be used as a starting point for building a vascularized breast tumor on-chip. The presence of breast vasculature interacting with and surrounding tumor spheroids, whether from established cell lines or from patient biopsies, could provide insightful information about the drug delivery process and its impact on the microenvironment.

## 5. Experimental Section/Methods

### Cell Culture

Commercially available Human Mammary Vascular Endothelial cells HMVEC were purchased from Innoprot (Cat # P10892) and were cultivated in endothelial media (VascuLife, Lifeline cell systems) with 5% Fetal Bovine Serum (FBS) on 30 μg/ml human fibronectin (Sigma) coated T-75 flasks. Primary Human Mammary Fibroblasts were purchased from Innoprot (Cat # P10893) and were cultured in Fibrolife media (Lifeline cell systems) on 50 μg/ml rat tail collagen I (Merck) coated T-75 flasks. All cells were used between passages 3 and 5. Human umbilical vein endothelial cells (HUVEC) were purchased from Lonza and transduced with viral particles generated in the lab by standard lentiviral particles production using the 3rd generation lentiviral expression vector pLV-mCherry (pLV-mCherry was a gift from Pantelis Tsoulfas) and cultured in endothelial media VascuLife on 50 μg/ml rat tail collagen I coated flasks. mCherry-HUVEC were used between passages 6 and 9. For all the cell lines used, the dissociations were carried out using TrypLE Express (Gibco), and growth media was completely refreshed every other day in the 2D cultures.

### Device Fabrication

As previously published [37], devices were fabricated using PDMS (SYLGARD™ 184 Silicone Elastomer Kit, Dow). The elastomer and cross-linker were mixed in a 10:1 ratio as per the manufacturer’s recommendation. Once mixed the PDMS is poured in a pre-fabricated multi-device negative mold and degassed using a vacuum desiccator. PDMS is then cured overnight at 60°C and then each device is cut and punched. Each PDMS device is air-plasma bonded (Harrick systems) to clean glass slides. The devices are coated with 30 μg/ml human fibronectin (Sigma) for 1 hour at 37 °C and after the devices were placed in the 60°C oven for 24 hours to return to its native hydrophobic state.

### Microvasculature Formation

Fibrinogen derived from bovine plasma (Sigma) was reconstituted in phosphate-buffered saline (PBS) to a working concentration of 6 mg/ml before use. Thrombin (Sigma) was diluted to a 4 U/ml working solution in cold VascuLife medium. HMVEC are stained with CellTracker™ Green CMFDA Dye (ThermoFisher) before the seeding to visualize vessel structures. The HMVEC and HMF were cultivated for two passages and when the confluence was reached they were dissociated and re-suspended at 12M cells/mL and 2.4 M cells/mL (5 to 1 ratio) in 4U thrombin. As previously described [37], the cells in thrombin were mixed with 50 %v/v of fibrinogen to make up final concentrations of 6 M cells/mL and 1.2 million cells/mL in 3mg/mL fibrin gel. The gel-cells mixture is inserted in the central channel of the device and to allow the polymerization of fibrin the devices were left at 37°C for 10-20 minutes. VascuLife with 5% FBS media was used to fill the media channels. After one day from the seeding, 3D printed custom-made reservoirs are inserted and used to generate an intermittent interstitial flow. For the 3 mm H2O pressure the media in the reservoir chamber is respectively 600 μL and 240 μL, while for the condition 0 mm H2O pressure in both reservoir chambers the media volume is 420 μL, refreshed daily. Microvessels were grown over 4 days prior to any treatments or permeability measurements.

### Permeability and morphological measurement

Endothelial barrier function was evaluated as a function of solute permeability, as previously discussed. [1, 36, 39] A 0.1 mg mL^−1^ solution of 70kDa TexasRed labeled dextran (ThermoFisher) in complete growth media (Vasculife) was perfused through the microvessels by a generated pressure drop across the gel. Both media channels were aspirated and then 40uL of dextran solution was first added to one media channel, to allow flow across the gel. This was then stabilized with the addition of an equal volume dextran solution to the opposing media channel to stop convective flow (∼30 seconds later.). All images were acquired on a Stellaris 8 confocal microscope using LAS X software (Leica). Intervals were set to 3 minutes between z-stacks. After 1 min, time-lapse (3 × 3-min intervals) confocal *z*-stack images were acquired at a 5 μm step size and ≈20–25 slices. Analysis was done as previously described in [36]. A maximum projection of the FITC-dextran channels at t = 0 and were used to quantify the morphology of the microvascular networks (for HMVEC-HMF and mCherryHUVEC-HMF co-cultures) as described in [36]. The morphological quantifications were then normalized to the size of the imaging region, as these measurements were performed on FITC and Texas red-dextran channels.

### 17β -estradiol and Progesterone Treatment

To emulate the menstrual cycle, the cells and the microvasculature have been treated with 17β -estradiol Bioreagent, G-irradiated (Merck, E2257-1MG), and with progesterone (Merck, P8783-5G) both reconstituted in absolute ethanol. Fixed concentrations from each phase of the menstrual cycle were determined from published plasma measurements [66, 71, 72]. Hormonal concentrations were as follows: follicular phase (0.3 nM E2,1.5 nM P4), ovulation (1.4 nM E2, 6 nM P4), and luteal phase (0.7 nM E2, 45 nM P4) and period (not treated).

### Cell extraction from microfluidic devices

The cell-gel channel of the devices was cut using a safety razor blade (VWR, Cat # 700-0418). The gels were digested in a solution of Accutase (Gibco) and 50 FU/ml Nattokinase (Japan Bioscience Ltd) for 15-20 min at 37°C. Afterwards, the content was homogenized and centrifuged. The remaining pellet was resuspended in 1mL of RNase-free water and stored at -70°C.

### RNA extraction and reverse transcription in cDNA

RNA has been extracted using the RNeasy Mini Kit (Qiagen, Cat # 74134) and performed according to the manufacturer’s handbook. The RNA samples have been diluted using 20 μL RNase-free water. The isolated RNA was converted to complementary cDNA using the High-Capacity RNA-to-cDNA™ Kit (Applied Biosystems, Cat # 4387406). Depending on the targeted cDNA concentration, the RNA volume for the reverse transcriptase was determined. The cDNA synthesis was performed in a thermocycler according to the manufacturer’s handbook. After reverse transcription, cDNA samples were stored at – 20°C until usage.

### RT PCR

To investigate the gene expression of the estrogen and progesterone receptors TaqMan real-time PCR was performed on 50 ng of cDNA per samples. The assay was performed using TaqMan Fast Advanced Master Mix (ThermoScientific, 4444556). The following primers were used to analyze the ER1 gene (ThermoScientific, Hs01046816), ER2 gene (ThermoScientific, Hs01100353), and PGR (ThermoScientific, Hs01556702). The assay has been performed in a 96-well plate (LightCycler® 480 Multiwell Plate 96, white, Roche) on the LightCycler ® 480 II (Roche). The PCR program includes a pre-incubation step at 50°C for 2 min and 95°C for 2 min, followed by 40 cycles of 1 second at 95°C and 20 seconds at 60°C for the amplification step. The gene expression data were normalized to the mean of two housekeeping genes ACTB (ThermoScientific, Hs01060665), TUB1A (ThermoScientific, Hs03045184), and tested in triplicates. The experimental data were analyzed with the 2-ΔCt method.

### Flow Cytometry

Characterization of HMVEC and HMF was done by flow cytometry. The HMVEC have been stained for endothelial markers: FITC Mouse anti-Human CD144 (BD Bioscience 560411, 1:50), PE-CF594 Mouse Anti-Human CD31 (BD Bioscience 563652, 1:100), PerCP-Cy™5.5 Mouse anti-Human CD105 (BD Bioscience 560819, 1:100). The HMF have been stained for the following fibroblast markers: Anti-S100A4 antibody Alexa647 (Abcam ab196168, 1:2000), Alpha-Smooth Muscle Actin Monoclonal Antibody (1A4), Alexa Fluor 488, (eBioscience 53-9760-82, 1:500), Anti-Vimentin Antibody (APC-Cy7) (Abcore AC12-0201-05, 1:500). Single cells were then stained with these markers for 2 hours at 4 °C, washed with PBS, analyzed on a BD LSR Fortessa and later processed using FlowJo v.10.8.1 software. The stromal cell population and the endothelial cells were gated using the positive cells to the antibodies and by the exclusion of the unstained cells cluster.

### Immunofluorescence staining

Fixation of cells and or devices were performed using 4% paraformaldehyde for 20 minutes prior to washing with PBS and subsequent solubilization using 0.1% Triton-X (10 minutes). To stain microvessels within a device, a pressure gradient was applied across the gel for all staining and wash steps. Samples were then incubated in applicable blocking buffer, PBS + BSA + serum of the secondary antibody, for more than 1 hour for 2D samples or overnight for the staining for the microvasculature. Primary antibodies were diluted in wash buffer (0.5% BSA in PBS) and were added to the samples and incubated overnight at 4°C. The primary antibodies used in the experiments are: Recombinant Anti-Estrogen Receptor alpha (phospho S118) antibody (Abcam ab32396, 1:100), Anti-Progesterone Receptor (phospho S190) antibody (Abcam, ab131110 1:100), Anti-CD31 antibody (Abcam, ab187377, 1:500) Recombinant Anti-S100A4 antibody (Abcam ab124805 1:500) and DAPI (FisherScientific, D3571) for the nuclei counterstaining. After overnight incubation, samples were washed with wash buffer and incubated with the appropriate secondary antibodies and counterstains (>2 h). Samples were rinsed with PBS and either imaged immediately or mounted on coverslips (for 2D samples) using Fluoromount-G (Invitrogen) and stored at 4°C before imaging.

### Cytokine array

For 3D cytokine analysis, supernatants were collected from 2D culture of HMVEC and HMF pooled from n=5 devices on day 4. We employed a human angiogenesis array (Abcam, ab134000) according to the manufacturer’s instructions. The relative expression of cytokines (measured by chemiluminescence intensity) was compared between all groups, corrected to the negative controls on each array, and normalized to the positive controls, using the monoculture as the reference array. The blots were visualized and imaged (1-minute exposure) using the Fusion FX Spectra (Vilber, France).

### ELISA assay

Quantification VEGF, bFGF, PLGF, and EGF in the media collected from each device (N=3 with 3 devices per experiment) was performed respectively with ELISA assay Human VEGF Quantikine ELISA Kit (R&D Systems, DVE00); Human FGF basic/FGF2/bFGF Quantikine Kit (R&D Systems, DFB50); Human PlGF Quantikine ELISA Kit (R&D Systems, DPG00) and Human EGF Quantikine ELISA Kit (R&D Systems, DEG00) according to manufacturer’s instructions. For this experiment, the supernatant was taken from the devices for the different hormone conditions on day 2 to exclude any influence of other processes involved in vessel regression, which could occur on day 4.

### MMP1 expression

Using DuoSet ELISA KIT (R&D systems, DY901B), serum MMP-1 levels were determined for microvessels on day 1 and 4 per the manufacturer’s protocol. Media samples were pooled from n=5 devices, immediately placed on ice, and frozen down at -70°C until assays were run. Samples were thawed on ice before the assay, while the other reagents were brought to room temperature before use (per manufacturer’s instructions). An appropriate sample dilution of 1:10 was determined to fit within the range of the MMP1 standard curve, using microvessel growth media as media control. Concentrations were measured using a plate reader at 450 nm (and a wavelength correction at 590 nm). Absorbance values were compared to provided standards, accounting for the sample’s dilution factor.

### Proliferation assay

To measure the proliferation of the cells in response to hormone treatment Click-iT Plus EdU Imaging Kits Protocol (Invitrogen,c10639) has been used according to the manufacturer’s instructions. Briefly, after 4 days of treatment with hormones, the samples are stained with EdU labeling solution (final concentration of 10 µM) for 2 hours. Then the cells are fixed with 4% PFA and permeabilized with (0.5% Triton® X-100 in PBS per 20 min). Later the cells are treated with Click-iT® Plus reaction cocktail for 30 minutes and then the nuclei are stained with Hoechst® 33342 for 30 minutes. The images (4 areas per well) were acquired with Thunder Imager Live Cell & 3D assay and using LAS X software (Leica) using the 10x objective.

### Statistical analysis

Statistical significance was analyzed using OriginPro v.9.85. For the samples that do not reject normality the one-way ANOVA was used to assess statistical significance across conditions at P < 0.05, and a post-hoc Tukey test was performed as a means comparison, where differences at p < 0.05 were taken as significant (*,p < 0.01 **, p < 0.001 ***, p < 0.0001 ****).The samples that reject normality the Kruskal-Wallis ANOVA was used to assess statistical significance across conditions.

### Computational fluid dynamics (CFD) simulations

CFD simulations were used to approximate the flow-induced shear stresses within the gel chamber of the microfluidic device. A CAD design of the complete microfluidic device including gel chamber that was sandwiched between fluid channels (.dxf file) was imported to COMSOL Multiphysics software (Version 6.0). This served as the geometrical input for the CFD simulation. Boundary conditions included the applied pressure gradient across the gel chamber (top to bottom) of the chip as mentioned in Figure S7 and no-slip conditions were applied to the side walls. Inlet pressure is set at 0.3 mbar (30 Pa or 3 mm H2O) and outlet at 0 mbar [73]. For this study, fluid properties of cell culture media were modeled as Newtonian fluid resembling water with a density of 998.2 kg/m^3^ and dynamic viscosity 9.4×10^−4^ Pa.s. Gel properties were modeled as having a density of 985 kg/m^3^and dynamic viscosity of 10^−4^1×10^−2^ Pa.s. Moreover, the porosity and permeability of the gel used were 0.3 and 1E^-15^ 1×10^−13^m^2^, respectively. To predict the flow-induced shear stresses and velocities profile within the gel chamber of a microfluidic chip, the Brinkman equation physics (flow through porous media) of COMSOL was used with physics-defined extremely fine mesh conditions.

## Supporting information

Supporting informations

## Acknowledgements

All Authors acknowledge the funding support from EMBL. We would like to acknowledge the Flow Cytometry Unit of the Center for Genomic Regulation for consultation in data acquisition and analysis, Joana Rafaela Mendonca da Silva for the help with data analysis.

