## Supporting informations for "Mammary microvessels are sensitive to menstrual cycle sex hormones"

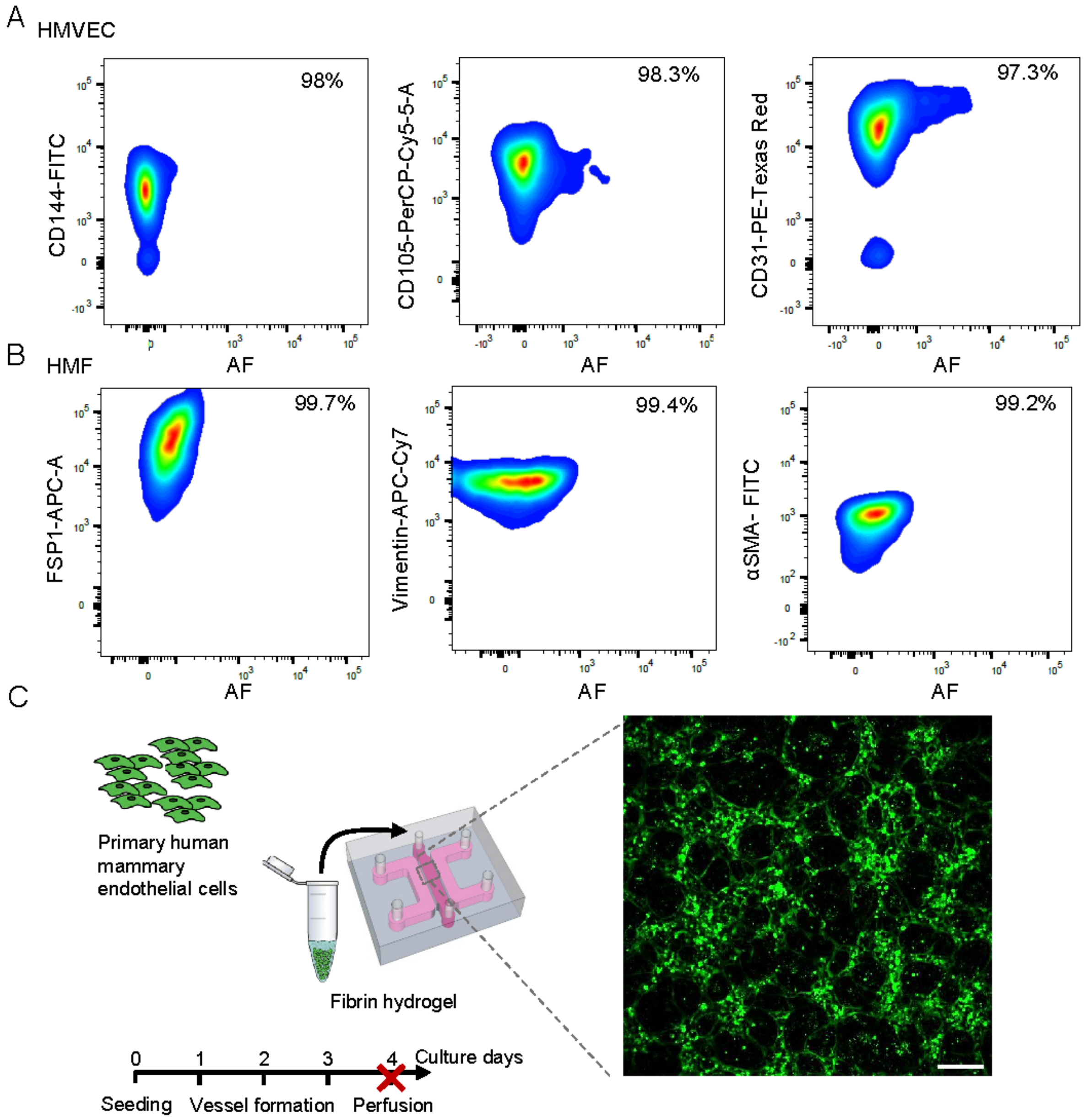


**Figure S1.** A) Flow cytometry characterization for HMVEC is shown for the following endothelial markers: CD144, CD105 and CD31. N=2 biological repeats. B) Flow cytometry characterization for HMF is shown for the following fibroblasts markers: FSP1, Vimentin and αSMA. AF = autofluorescence. C) (Left) Graphical representation of the procedure used to generate mammary microvessels with HMVEC only (absence of fibroblasts). Representative confocal maximum projection of HMVEC in the system shown with CellTracker™ at day4. Scale bar is 200 μm.

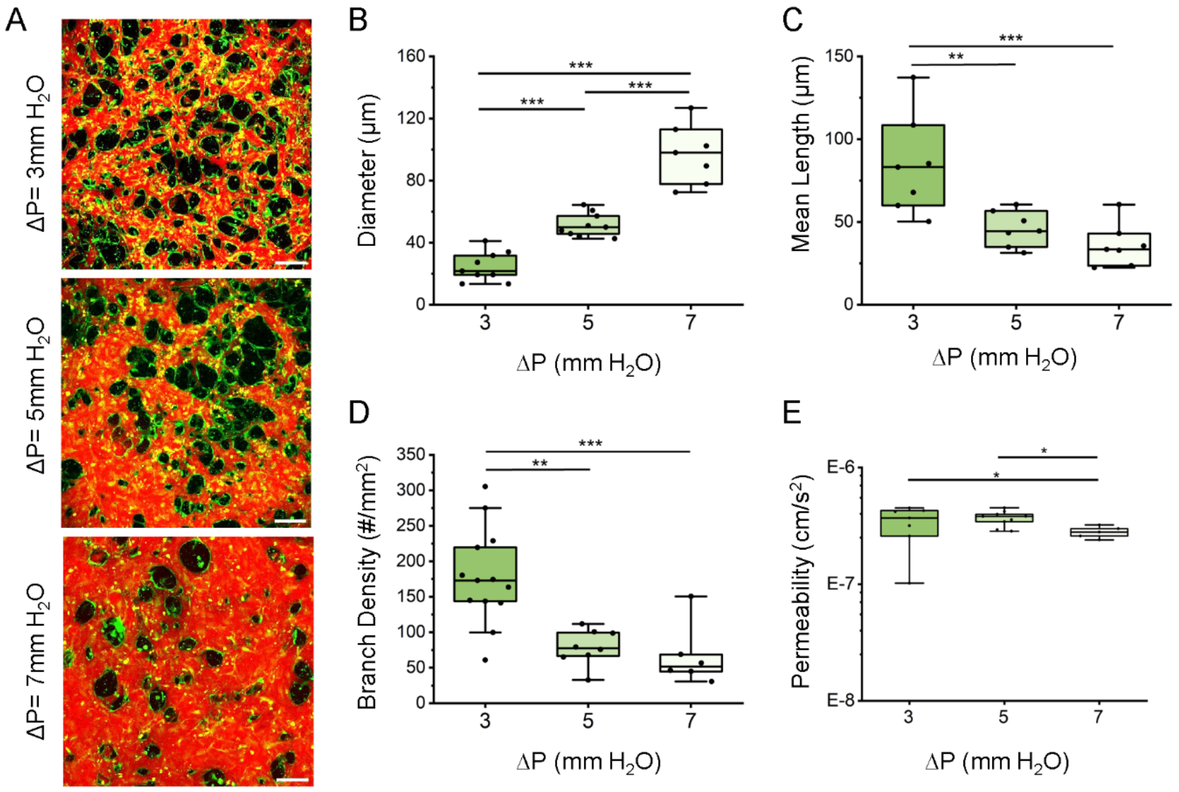


**Figure S2.** A) Confocal maximum projection images highlighting microvessels formed applying different pressure gradients –which induces flow at: ΔP = 3, 5 and 7 mm $H_2O$. HMVEC are shown with CellTracker™ green and microvessels are perfused with 70 kDa dextran (red) at day 4 to assess vessel permeability. Scale bar is 200 µm. Comparison of the microvessels across culture conditions for B) vessel effective diameter, C) mean vessel length, D) branch density and E) vessel permeability, for N=2 biological repeats. Box plots demonstrate median, percentile 25-75 quartile (box edge) and 10-90 (outer whiskers). Significance is shown by $*p < 0.05$, $**p < 0.01$, $***p < 0.001$ using one-way ANOVA with Tukey means comparison test.

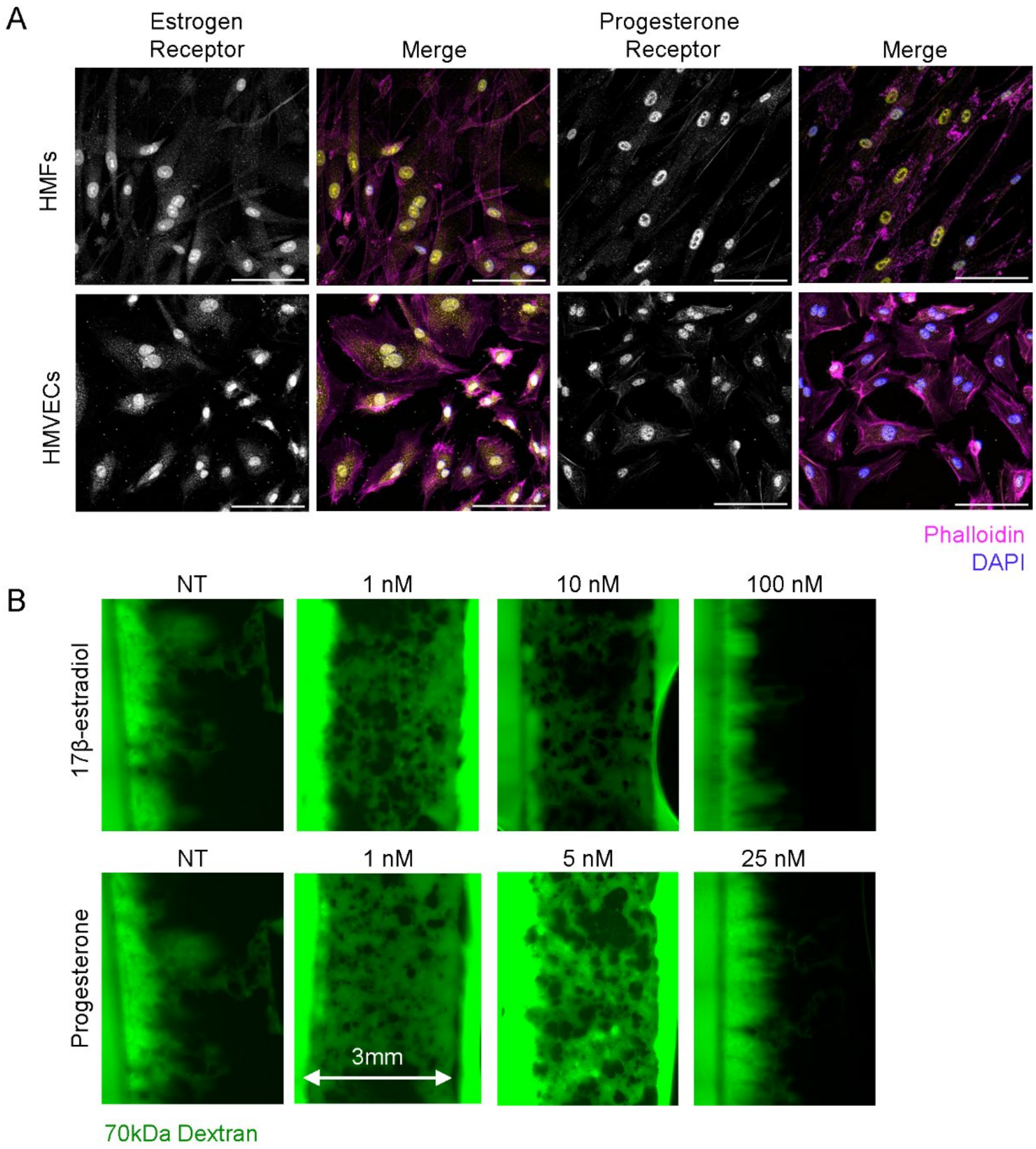


**Figure S3 .** A) Immunofluorescent images of HMF and HMVEC stained with estrogen and progesterone receptors. Scale bar is 200 µm. In the merge, yellow are the receptors, magenta is phalloidin and in blue DNA. B) Overview image of the HMVEC–HMF co-cultured microvessels treated with 1, 10 and 100 nM of 17β-estradiol and 1, 5 and 25 nM of Progesterone. Non-treated controls (NT) correspond to samples without hormones. The microvessels are perfused with 70 kDa FITC dextran.

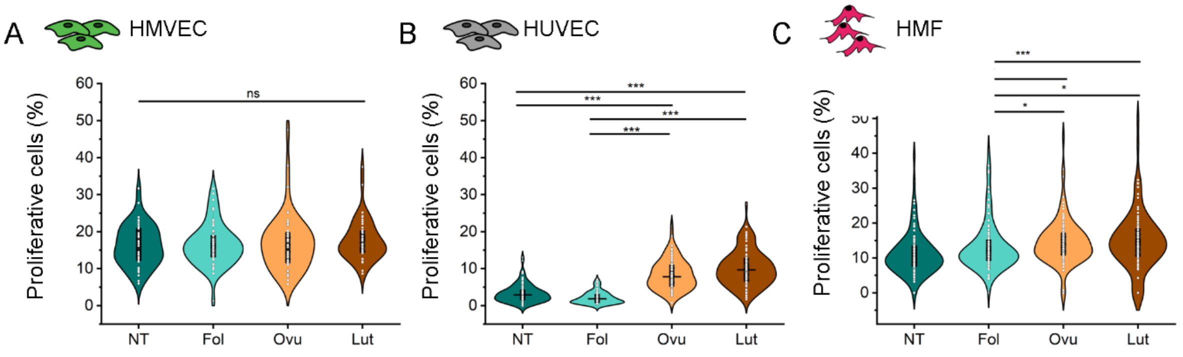


**Figure S4 .** Shown are the percentage of proliferative cells for A) HMVEC, B) HMF and C) HUVEC, cultured in 2D with the different hormonal treatments (non-treated, NT; Follicular, Fol; Ovulation, Ovu; Luteal, Lut). Significance is shown by *$p < 0.05$, **$p < 0.01$, ***$p < 0.001$ using one-way ANOVA with Tukey means comparison test. Violin plots demonstrate median, SD (outer whiskers) and SE (box edge). N=3 biological repeats.

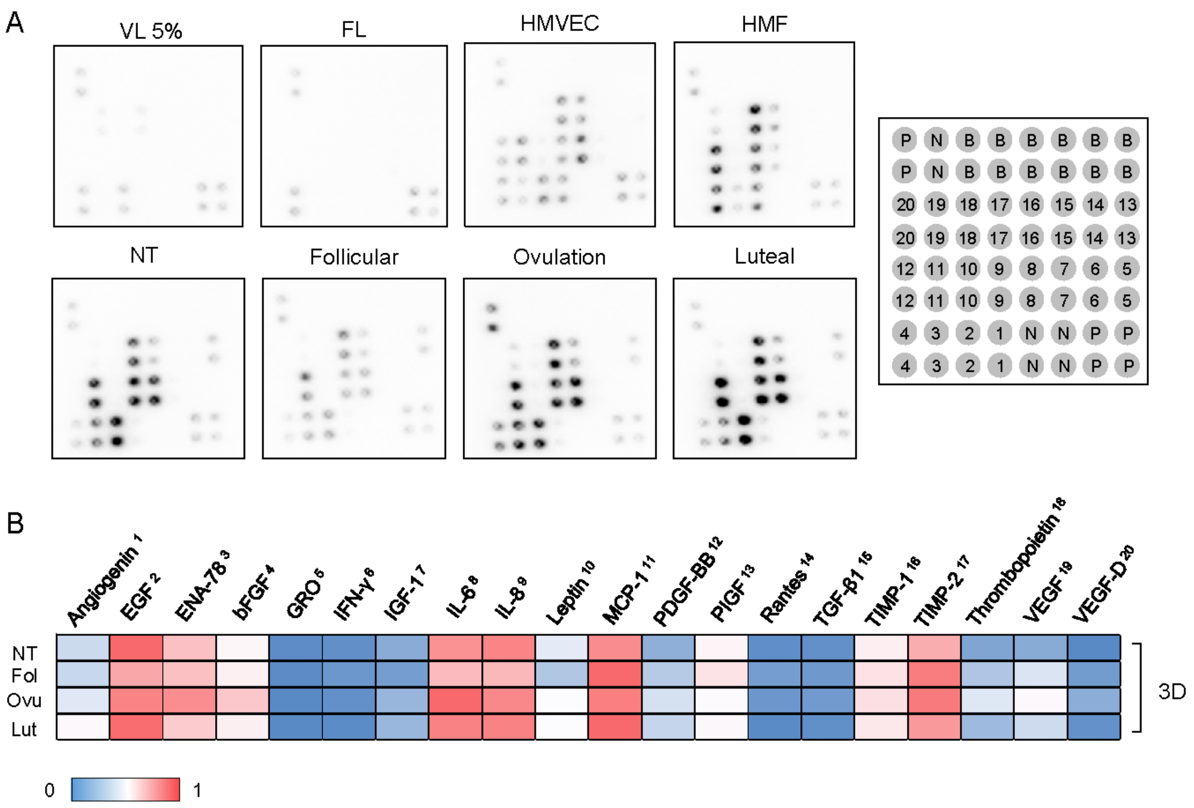


**Figure S5 .** A) Raw data showing cytokine array from pooled n=4 supernatant collected at day 4 from vessels treated with the different hormonal treatment conditions (non-treated, NT; Follicular, Fol; Ovulation, Ovu; Luteal, Lut). B) Semi-quantitative analysis shows intensity differences between conditions.

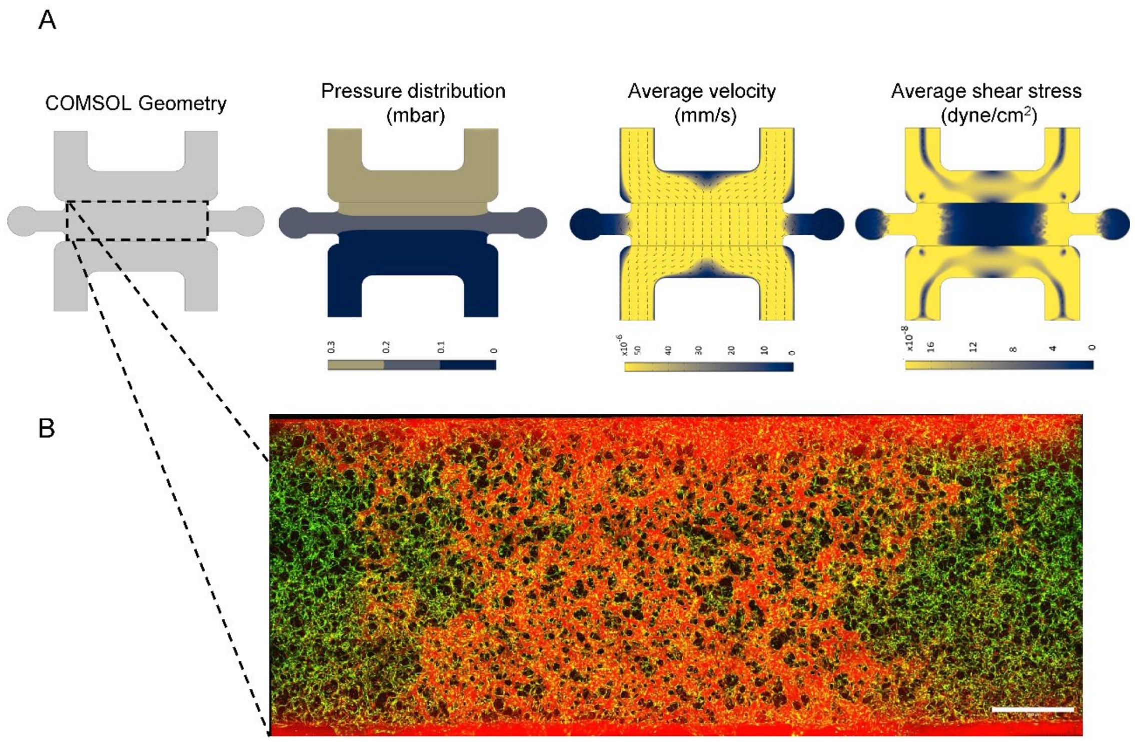


**Figure S6 .** A) The microfluidic design was imported into COMSOL multiphysics software, where an applied pressure gradient across the gel channel is used to predict velocities and shear stresses within the gel and media channels, respectively. B) Confocal maximum projection images of an entire device showing mammary microvessels at day 4. HMVEC are shown using CellTracker™ green and perfused vessels with 70 kDa dextran (red). Scale bar is 1000 µm.

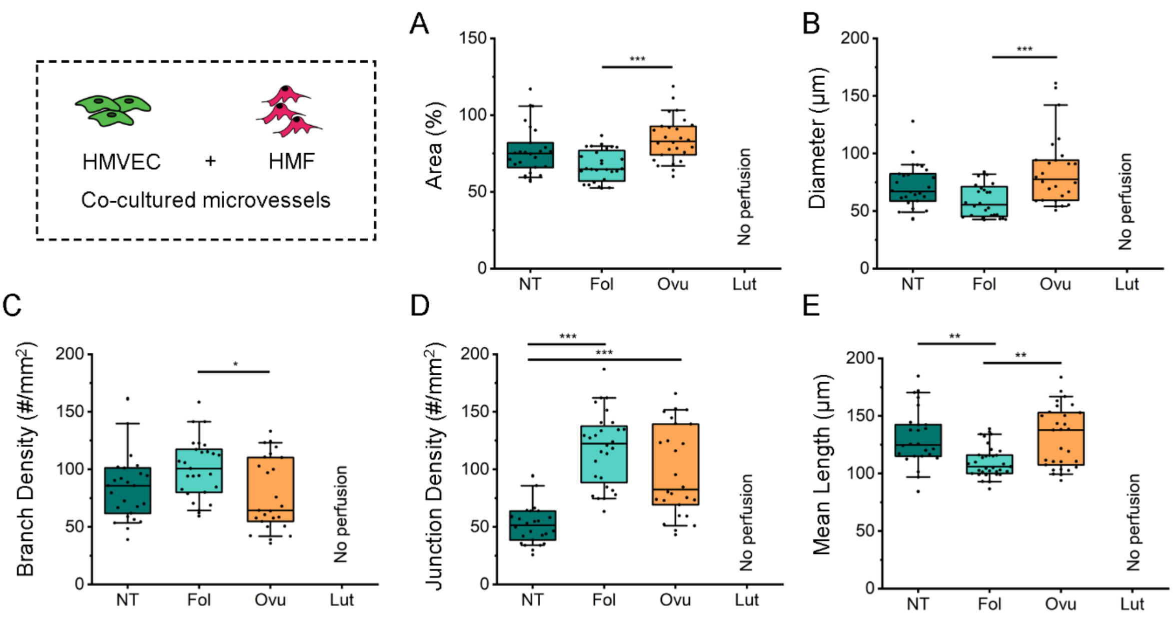


**Figure S7 .** Morphological comparison of the breast microvessels with the different hormonal treatments (non-treated, NT; Follicular, Fol; Ovulation, Ovu; Luteal, Lut). Non-normalized data is shown for A) vessel area coverage, B) effective diameter C) branches density, D) junction density and E) vessel length. Box plots demonstrate median, percentile 25-75 quartile (box edge) and 10-90 (outer whiskers) for N=5 biological repeats. Significance is shown by *$p < 0.05$, **$p < 0.01$, ***$p < 0.001$ using Kruskal-Wallis ANOVA test.

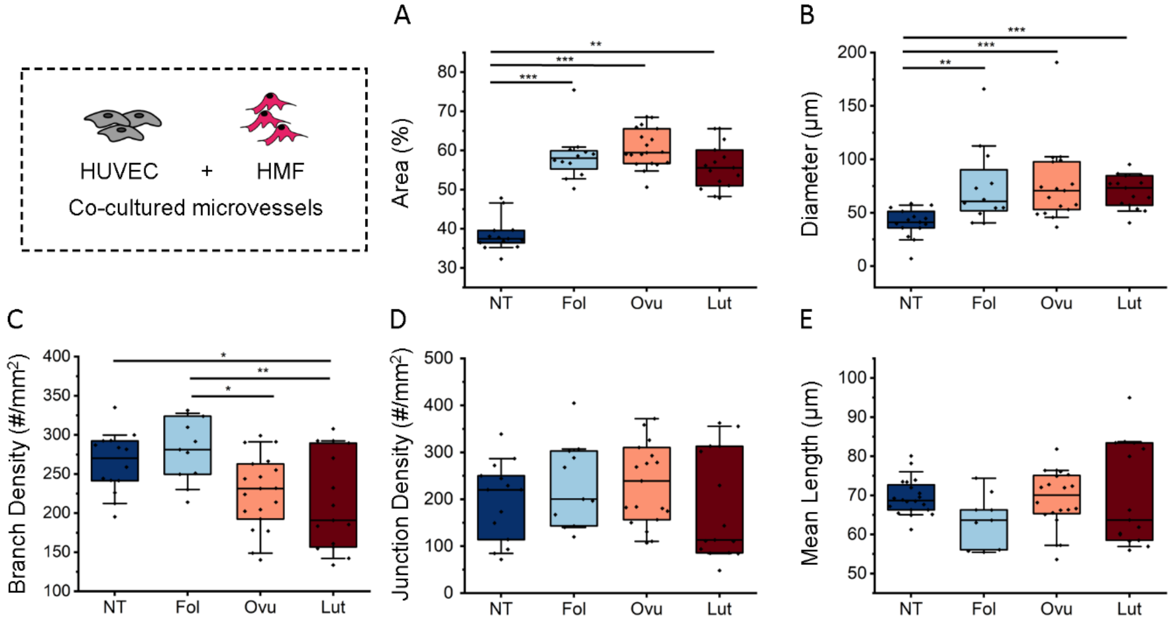


**Figure S8 .** Morphological comparison is shown for HUVEC-HMF co-cultured microvessels exposed to different hormonal treatments (non-treated, NT; Follicular, Fol; Ovulation, Ovu; Luteal, Lut). Shown is A) vessel area coverage, B) effective diameter, C) branches density, D) junction density and E) vessel length. Box plots demonstrate median, percentile 25-75 quartile (box edge) and 10-90 (outer whiskers) for N=3 biological repeats. Significance is shown by *$p < 0.05$, **$p < 0.01$, ***$p < 0.001$ with one-way ANOVA and Tukey means comparison test for data following normality, or if normality is rejected using Kruskal-Wallis ANOVA test.
